# Bridging Biomedical Atlas Ecosystem: Cross-Atlas Alignment And Scalable Tissue Specimen Registration

**DOI:** 10.64898/2026.08.13.744704

**Authors:** Yashvardhan Jain, Bhargav Desai, Danial Qaurooni, Antara Bhavsar, Peter Kienle, Alison M. Pouch, Kathleen O’Neill, Siddharth Apte, Bruce W. Herr, Stephen A. Fisher, Katy Börner

**Affiliations:** Department of Intelligent Systems Engineering, Luddy School of Informatics, Computing, and Engineering, Indiana University, Bloomington, IN 47408, USA; Departments of Radiology and Bioengineering, University of Pennsylvania, Philadelphia PA 19104, USA; Department of Obstetrics and Gynecology, University of Pennsylvania, Philadelphia PA 19104; Department of Biology, University of Pennsylvania, Philadelphia PA 19143, USA

## Abstract

Over the last five years, over 13,000 tissue datasets with 200+ million cells from 20 consortia have been spatially registered into the Human Reference Atlas (HRA) common coordinate framework (CCF). The shared 3D spatial and semantic reference system enables exploration of datasets in the context of all other data across organs, assay types, and spatial scales.

However, manual registration of individual samples remains resource intensive, posing feasibility challenges exacerbated by the proliferation of samples, assays, and atlasing efforts. This paper presents two approaches to scale up HRA construction: (1) projecting data across biomedical reference atlas systems and (2) using millitomes to bulk register tissue blocks into a reference organ. Both methods use the **A**uto**M**ated **A**lignment and **P**rojection (AMAP) pipeline to align 3D mesh models using point cloud registration. We demonstrate the evolving HRA-aligned atlas ecosystem for 6 models from the SPARC Program (heart), Gut Cell Atlas (large intestine), 500-subject consensus kidneys, and the Julich Brain Atlas. Additionally, we used AMAP to project 7 millitome models across 5 organs onto the HRA ecosystem, integrating 300+ tissue extraction sites. AMAP enables scalable tissue registration of data across atlas ecosystems enabling the construction of detailed reference maps of the human body.

## Introduction

The Human Reference Atlas (HRA)^1,2^ is a multiscale, multimodal, three-dimensional (3D) atlas of the anatomical structures, cells, and biomarker expression values in the healthy human body.

Funded by the National Institutes of Health (NIH) and with expert support by the Human Cell Atlas (HCA) effort^3,4^, the HRA brings together 2D and 3D tissue specimen data across organs, assay types, and data providers, with contributions from experts across 25+ international consortia, including the Human BioMolecular Atlas Program (HuBMAP)^5,6^, the Cellular Senescence Network (SenNet)^7,8^, and the Kidney Precision Medicine Project (KPMP)^9,10^.

To register tissue data into the male and female reference bodies, the HRA provides the Registration User Interface (RUI)^11^; tissue data can then be explored in the Exploration User Interface (EUI)^11^. Together, these two interfaces make it possible for subject matter experts to register tissue samples spatially and annotate them semantically. As of July 13, 2026, 54 tissue data providers have used the RUI to register 2,762 tissue blocks from 1,021 unique donors that are linked to 13,344 tissue datasets that can be explored at https://apps.humanatlas.io/eui.

Besides HuBMAP, SenNet, and KPMP, contributing consortia include among others the Stimulating Peripheral Activity to Relieve Conditions (SPARC) Program^12,13^, the GenitoUrinary Development Molecular Anatomy Project (GUDMAP)^14,15^, the Genotype-Tissue Expression (GTEx) Program^16,17^, the Human Cell Atlas (HCA)^3,4^, the Chan Zuckerberg Initiative CELLxGENE program^18^, BRAIN Initiative Cell Census Network Initiative (BICCN)^19,20^, and the Human Tumor Atlas Network (HTAN)^21^.

Going forward, the HRA must keep pace with the rapid proliferation of tissue samples, assay types, and parallel atlasing efforts. Repeated collaborations with tissue providers following standardized protocols need to scale beyond individual tissue block registration toward high-throughput processing pipelines. Simultaneously, projecting data across organ-specific atlases—such as the Julich Brain Atlas^22^ and the Helmsley Gut Cell Atlas^23,24^—enables their tissue data to be surfaced within the HRA EUI and vice versa, improving data findability, accessibility, interoperability, and reusability (FAIR). Together, these demands point to the need for an automated, scalable, cross-atlas spatial registration workflow.

This paper presents **A**uto**M**ated **A**lignment and **P**rojection (AMAP), a Python-based pipeline for point-cloud-based registration (transformation and projection) of 3D mesh organ models, enabling cross-model tissue block projection through whole-organ alignment. AMAP transforms organ models into point-cloud representations and applies a sequence of linear, affine, and deformation-based transforms to align source and target models, which enables two applications (see **Figure 1**):

1. **Cross-atlas alignment:** We use AMAP to align HRA 3D reference organ models with those from the SPARC heart^25^, Helmsley Gut Cell Atlas (GCA) large intestine^23^, VU500 consensus kidneys^26^, and Julich Brain Atlas^22^, projecting HRA-registered tissue blocks into each system.
2. **Millitome-based registration:** To scale tissue registrations into the HRA, we collaborated with data providers to create millitome-based 3D models according to their tissue sectioning protocols^27–29^. We showcase millitome construction and subsequent tissue block registration for kidney, pancreas, and female reproductive organs across multiple HuBMAP tissue providers.

**Figure 1.**
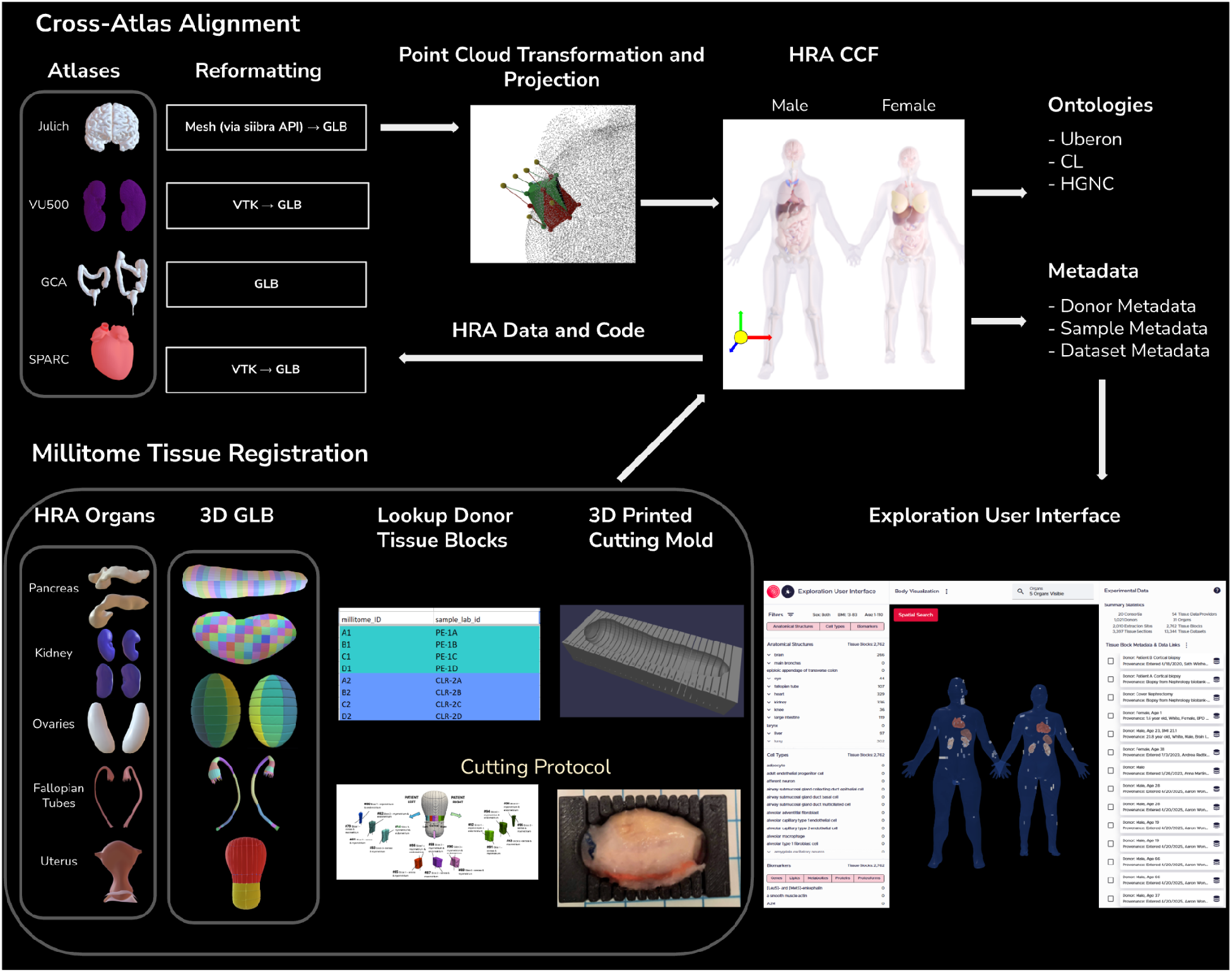
Workflow Overview. **Top panel.** An overview of the Cross-Atlas Alignment user needs scenario. Four different atlases (Julich Brain Atlas, VU500 Consensus Kidney Atlas, Gut Cell Atlas Large Intestine, and SPARC Heart Scaffold) are reformatted into GLB formatted 3D models, which are then converted into point clouds for transformation and projection between the HRA 3D Reference Models using AMAP pipeline. The extraction sites and tissue blocks (and associated HRA CCF ontologies and metadata) can then be pulled via HRA APIs and projected into relevant atlases. **Bottom Panel.** An overview of the Millitome Tissue Registration user needs scenario. The HRA 3D Reference Models for five organs (pancreas, kidneys, ovaries, fallopian tubes, and uterus) are shown along with 3D millitome models (virtual GLB files); each virtual model has a lookup table associated with it. A millitome can also be implemented as a 3D printed cutting mold as well as surgeon instructions (or cutting protocol). The Exploration User Interface (EUI) shows the tissue blocks registered in the HRA CCF using either the Registration User Interface (RUI) or millitome-based registration. The same data can be projected into other atlas systems using the point cloud transformation and projection methods implemented and demonstrated in this paper.

**Figure 2.**
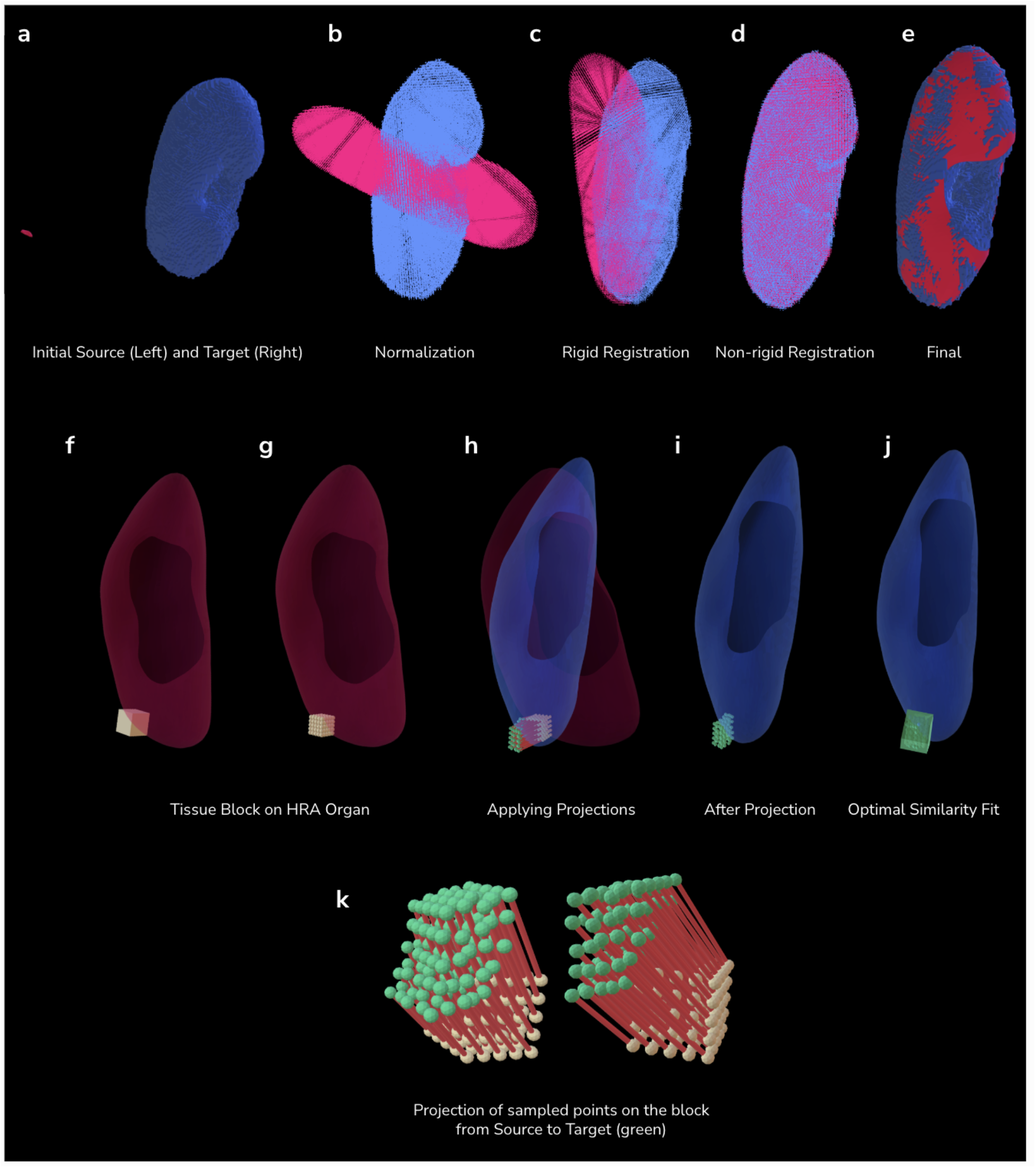
Overview of the AMAP whole organ transformation and tissue-block projection pipeline. **a.** Initial source and target organ meshes before transformation (the HRA organ in red was enlarged 100 times to make it visible); **b.** Normalized source and target geometries in a common frame; **c.** Rigid registration (transformation); **d.** Non-rigid registration (transformation, point cloud); **e.** Final transformed source in target space (mesh); **f.** Dummy tissue block that sticks out of kidney for visibility registered into the HRA source model; **g.** Dense sampling of the tissue block bounds for points to be propagated through the projections; **h.** Transformed block points (green) after projection of the tissue block (beige) shown with reference to the registered source (blue) and target anatomy (red); **i.** Projected dense samples after projection; **j.** Final projected tissue block obtained after optimal similarity fitting between the sampled points (beige) and the projected points (green); **k.** Transformation of the sampled points from the source block to their projected target positions by the deformation vector field (from the non-rigid transformation).

Together, AMAP and millitomes make it possible to register and exchange tissue datasets at scale and across atlas efforts, effectively creating a reference atlas ecosystem.

### Box 1. Key Terminology

- **AMAP**: This acronym refers to the process of aligning two mesh-based 3D models and projecting the underlying data between human atlas systems for data and code interoperability. In the Human Reference Atlas this is used to project registered tissue blocks from a specific source organ to a reference organ.
- **Alignment:** Process for transforming the geometry/shape of a source mesh (or point-cloud) model to a target mesh (or point-cloud) model using 3D point-cloud-based registration (linear, affine, and deformation-based transforms).
- **Projection:** Post-alignment process for computing coordinates of the underlying data (extraction sites) based on the computed transformations between the target model and the source model.
- **Spatial Registration:** The process of mapping tissue samples into reference anatomical bodies. It combines spatial alignment (positioning samples relative to the reference) with semantic annotation (linking sample regions to defined anatomical structures) to enable comparison and integration across datasets.
- **Tissue Block:** A sample or specimen derived from an organ or tissue including subsections thereof obtained from a donor that has a unique ID and links to donor, organ extraction site, processing, preservation and other metadata. The locations of tissue blocks are registered using the Registration User Interface.
- **Extraction Site**: Digital, 3D representation of a tissue block. If the Registration User Interface is used to register tissue, then each site has a unique ID; data on size, location, and rotation in 3D in relation to an Human Reference Atlas reference organ; a listing of all anatomical structures that the cuboid intersects with (bounding-box collision by default); and metadata on who registered it.
- **HRA Common Coordinate Framework (CCF):** As used in the Human Reference Atlas, this consists of ontologies and reference object libraries, computer software (e.g., user interfaces) and training materials that (1) enable biomedical experts to semantically annotate tissue samples and to precisely describe their locations in the human body (“registration”), (2) align multi-modal tissue data extracted from different individuals to a reference coordinate system (“mapping”), and to (3) provide tools for searching and browsing data at multiple levels from the whole body down to single cells (“exploration”).
- **Millitome:** Millitomes provide guidance on how to partition an organ, often a complete organ. Millitomes can include either a 3-dimensional mold that physically holds an organ for partitioning, or a drawing of an organ that is used as a guide by a surgeon when they partition an organ free hand, or both. Using a millitome ensures that tissue sample locations show correctly in the Exploration User Interface.

## Results

### User Needs

This paper presents two symbiotic approaches to scale up human reference atlas construction and usage: (1) cross atlas alignment and (2) using millitomes to bulk register hundreds of tissue blocks into a reference organ. Both are implemented using **A**uto**M**ated **A**lignment and **P**rojection (AMAP), a Python-based pipeline for point cloud transformation and projection.

#### Cross-Atlas Alignment

Over the last 10 years, there has been a proliferation of human atlas projects^6^. Many are organ specific, e.g., Allen^30^ and Julich^22^ for brain, KPMP^9,10^ and VU500^26^ for kidney, GCA^23^ for large intestine, see **Figure 1**, top left. The Human Reference Atlas (HRA)^2^ aims to cover vital organs initially and all organs ultimately. Data and code interoperability is highly desirable^31^ as organs are highly dependent on the functionality of other organs (e.g., the brain needs the lung for oxygen and intestine for nutrients) but difficult due to different data formats and coordinate systems used for the 3D reference scaffolds representing the anatomical structures of organs. For example, the SPARC heart and VU500 kidneys come in VTK format, Julich brain in a mesh via siibra-python API^32^, HRA in GLB format. The coordinate system origin (i.e., 0, 0, 0) for Julich brain and SPARC heart models (GLB files) is in the approximate geometric center while it is the bottom-left-back for the HRA. Cross-Atlas Alignment shown in the top part of **Figure 1** makes it possible to translate data across atlases:

(1) reformat all data into GLB format and (2) perform cross-atlas alignment using point cloud transformation and projection. This translation of the data enables the use of data from other atlases in the HRA (e.g., to explore it in EUI in the context of more than 10,000 other datasets across 30+ organs) and to use HRA data and code (e.g., the RUI for tissue registration) in other atlasing efforts.

#### Millitome Tissue Registration

Some atlas teams have access to entire organs, segment these into tens to hundreds of tissue blocks, and are interested to register all “in bulk” and not “one-by-one” via the RUI^11^ (https://humanatlas.io/registration-user-interface). Inspired by initial work^11^ by the HuBMAP tissue mapping center at Vanderbilt University, we developed millitomes to bulk register hundreds of tissue blocks into an HRA reference organ. A millitome defines a protocol for scaling up tissue registration by partitioning a human organ model and registering the resulting tissue blocks into extraction sites within the HRA three-dimensional (3D) reference objects. The protocol, e.g., by GTEx^33^, can be implemented in two ways: (i) using a 3D (printed) Cutting Mold or (ii) a Cutting Protocol used by surgeons to cut tissue blocks in a very systematic, reproducible manner. In both cases, 3D GLB files with pre-segmented virtual tissue blocks exist and need to be aligned to the HRA organs so tissue blocks that have experimental data appropriately located in the Exploration User Interface. Here, AMAP is used to align millitome models to HRA 3D reference models and all individual blocks within the millitome geometry are “registered” in the HRA and visualized in the EUI, thereby bypassing the “one-by-one” registration process of RUI. Lookup Donor Tissue Blocks CSV files are used to capture donor and other metadata. The process is shown in the lower part of **Figure 1**.

Subsequently, we present the general AMAP Point Cloud Transformation and Projection approach followed by details on Cross-Atlas Alignment and Millitome Tissue Registration.

### Point Cloud Transformation and Projection

**A**uto**M**ated **A**lignment and **P**rojection (AMAP) is a Python-based pipeline that computes linear, affine, and deformation-based transformations between two 3D models (source model and target model) using point-cloud-based registration algorithms. The goal is to align the source model to the target model as accurately as possible based on surface geometries.

While AMAP supports 3D mesh data formats such as STL and GLB as inputs, the mesh-based geometries are first converted into point clouds (see **Data Formats** in **Methods, and Figure 1 top panel**) for both source and target models before the alignment process. The alignment process is implemented as two distinct steps:

1. Rigid registration via Random Sample Consensus (RANSAC)^34^ and Iterative Closest Point (ICP; point-to-plane variant)^35^ produces an initial coarse alignment between source and target point clouds, resolving laterality differences—such as left-to-right kidney flips—and reconciling organs from different coordinate spaces.
2. Non-rigid registration using Bayesian Formulation of Coherent Point Drift (BCPD++)^36,37^ refines this alignment by estimating deformation parameters under a motion coherence prior, with manually tuned hyperparameters (see **Methods**) governing motion vector length and rigidity. The final outputs—a transformation matrix, a deformation vector field (DVF)—are serialized into a Python object (saved as a *\*.pickle.gz* file) for downstream use.

Once whole organ alignment is computed, projecting the underlying tissue blocks (i.e., their extraction sites) across source/target coordinate spaces is straightforward: the combined transformations describe the displacement of each source mesh vertex required to match the target, and these displacements are applied to the vertices of any tissue block (extraction site) registered within the source organ. Vertices falling between mesh resolution points are handled via nearest-neighbor interpolation. BCPD++ also generates probabilistic correspondences between point clouds, but for anatomically complex organs such as the brain or kidney, strict point-to-point correspondence is not guaranteed, particularly when source and target point clouds differ in density. The DVF therefore serves as the primary basis for projection.

The AMAP registration pipeline is designed to be accurate enough to approximately visually align two datasets. It is not designed for point perfect registration. For analysis relying on highly accurate registration between two models, a more accurate local registration should be computed, using AMAP registration as an initial starting point. Further implementation details are provided in **Methods**.

### Cross-Atlas Alignment

Atlas construction efforts across the biomedical community have produced CCFs of varying spatial dimensionality. AMAP is used to align these non-HRA models—the Gut Cell Atlas large intestine^23^, SPARC heart^25^, VU500 consensus kidneys^26^, and Julich Brain^22^—to their corresponding HRA counterparts (from the HRA 3D reference object library). Prior to alignment, data format conversions are performed as needed for compatibility with each reference system (see **Methods**).

Once alignment is computed, RUI-registered tissue blocks in the HRA are projected into non-HRA coordinate spaces based on the optimized set of transformations between the source (HRA) and target (non-HRA) models. This is useful in contexts where users prefer to explore tissue data within their own atlas reference system—for example, the Gut Cell Atlas encodes intestinal location as a percentage of organ length, mirroring clinical colonoscopy practice, while SPARC references support computational models that simulate a beating heart or peristaltic colon movement. **Figures 3-5** show HRA tissue blocks projected into the SPARC heart, Helmsley Gut Cell Atlas, VU500 kidney, and Julich Brain models. Each aligned model, including projected extraction sites, is exported as a GLB file with extraction sites as separate mesh geometries, accompanied by JSON-LD files containing block-level metadata from the HRA API. Since alignments can be computed bidirectionally, tissue data registered in non-HRA systems can similarly be projected back to the HRA and displayed in the EUI.

**Figure 3.**
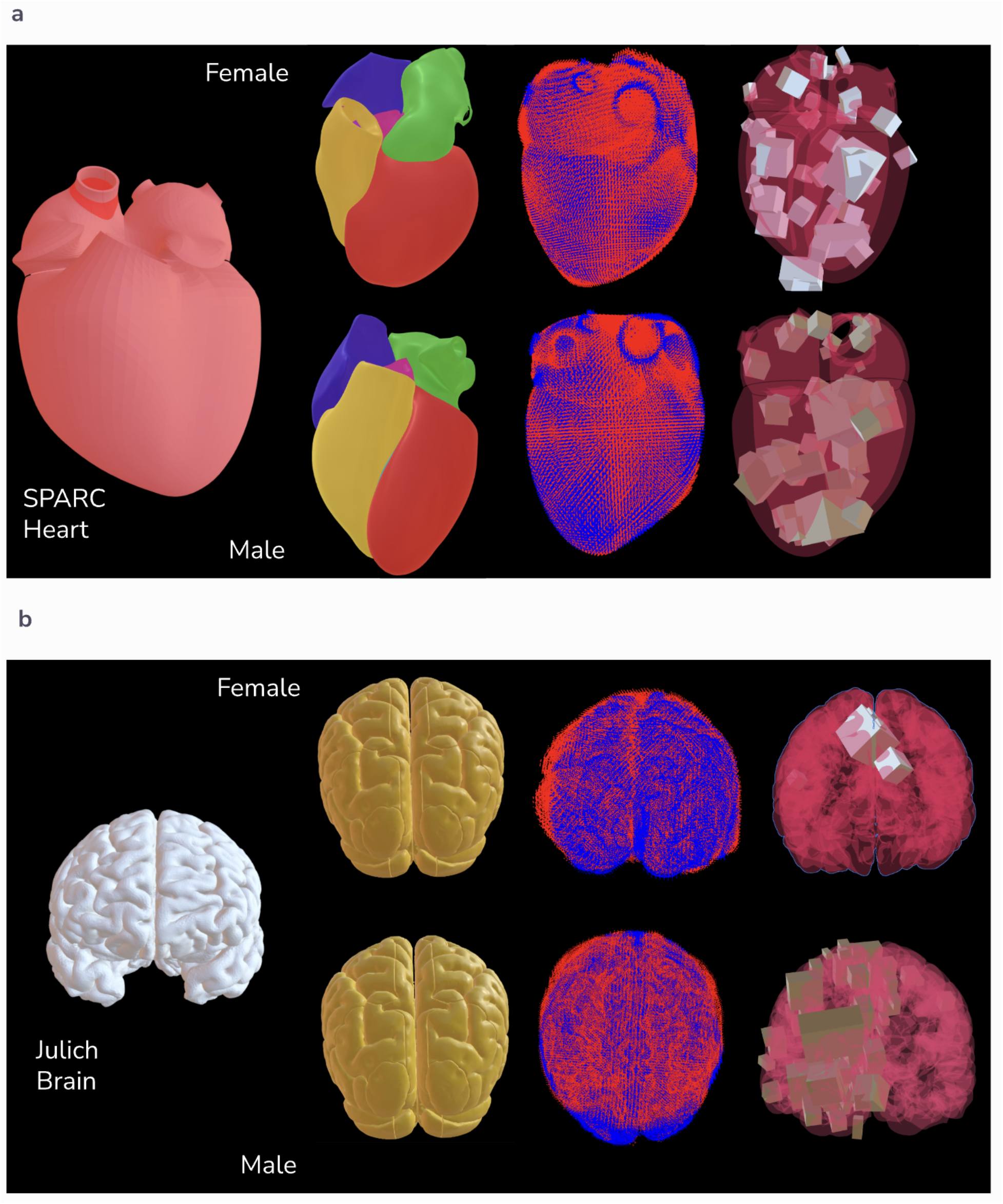
Cross-Atlas Alignment for SPARC Heart and Julich Brain. **a.** Shows the SPARC heart scaffold (3D GLB model; target model) and the HRA 3D Reference Models (male and female; source models), followed by the point cloud overlaps (after AMAP transformation; blue: target model, red: source model), finally showing the HRA tissue blocks (from EUI) projected into the SPARC heart model. **b.** Shows the Julich brain model (3D GLB model for *MNI_COLIN_27* template space; target model) and the HRA 3D Reference Models (male and female; source models), followed by the point cloud overlaps (after AMAP transformation; blue: target model, red: source model), finally showing the HRA tissue blocks (from EUI) projected into the Julich brain model.

**Figure 4.**
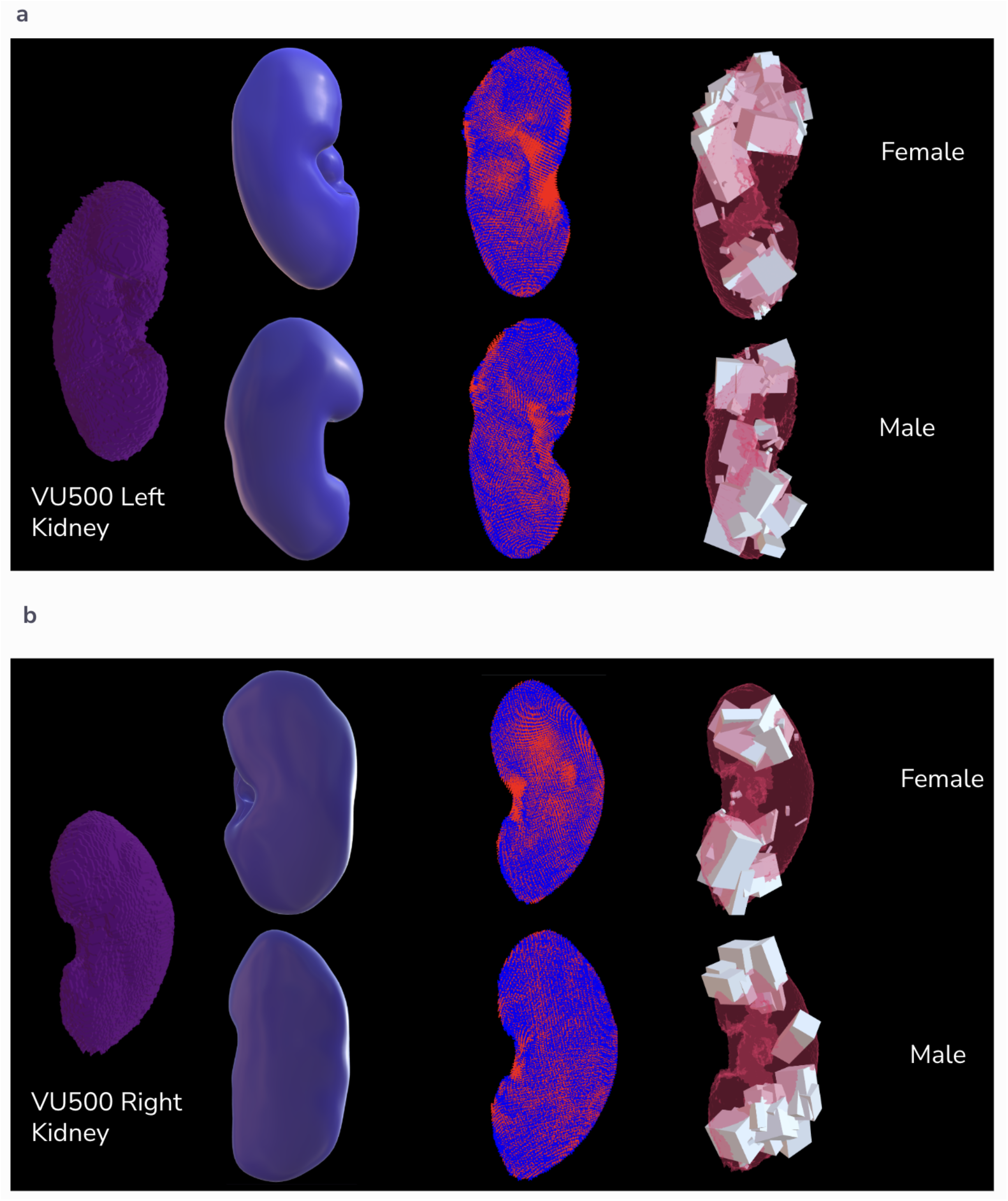
Cross-Atlas Alignment for VU500 Consensus Kidneys. **a.** Shows the VU500 Consensus left kidney (3D GLB model; target model) and the HRA 3D Reference Models (male left and female left; source models), followed by the point cloud overlaps (after AMAP transformation; blue: target model, red: source model), finally showing the HRA tissue blocks (from EUI) projected into the left consensus kidney. **b.** Shows the VU500 Consensus right kidney (3D GLB model; target model) and the HRA 3D Reference Models (male right and female right; source models), followed by the point cloud overlaps (after AMAP transformation; blue: target model, red: source model), finally showing the HRA tissue blocks (from EUI) projected into the right consensus kidney.

**Figure 5.**
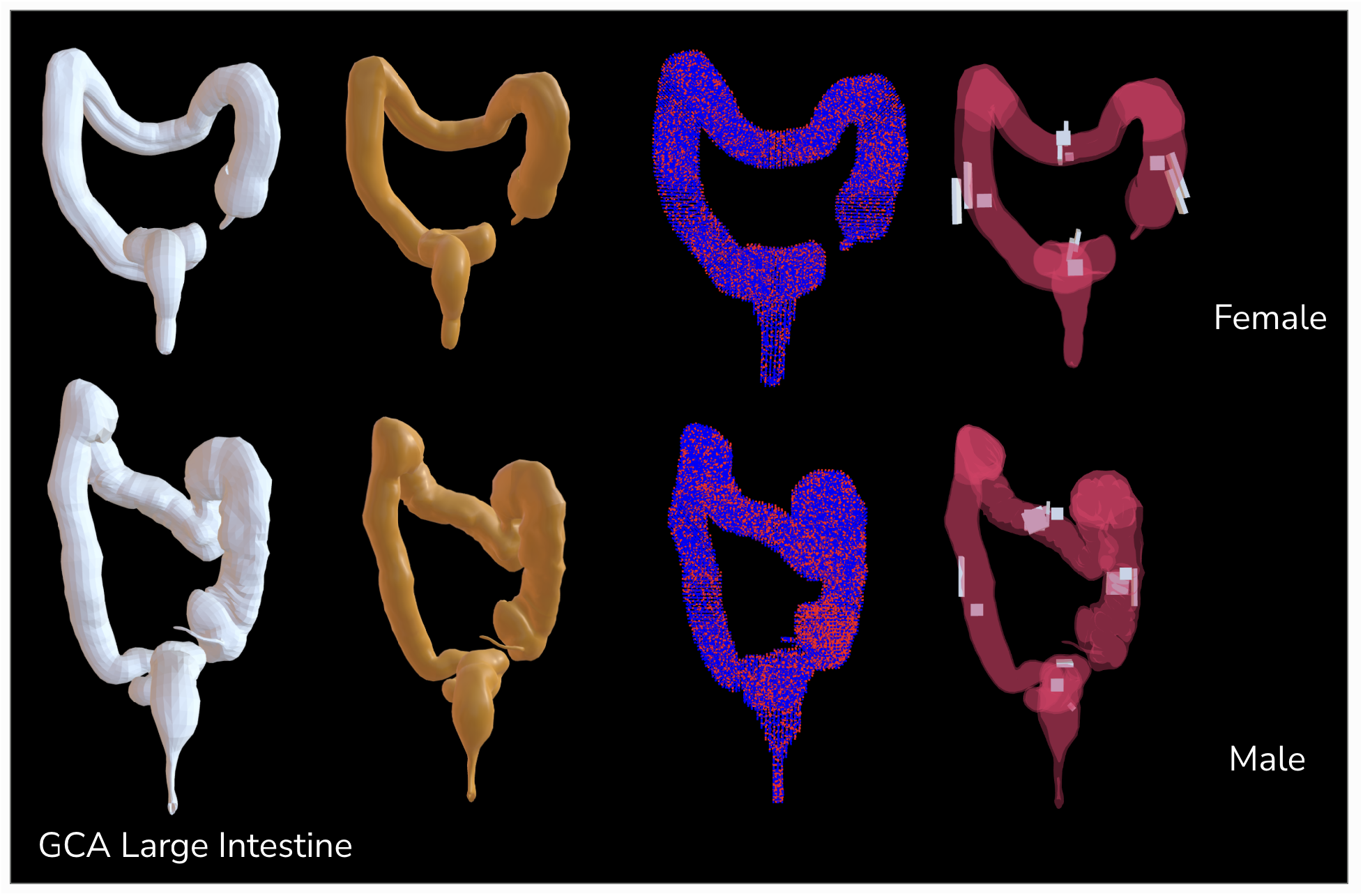
Cross-Atlas Alignment for Gut Cell Atlas (GCA) Large Intestines. Shows the female and male GCA large intestines (3D GLB models; target models) and the HRA 3D Reference Models (female and male; source models), followed by the point cloud overlaps (after AMAP transformation; blue: target model, red: source model), finally showing the HRA tissue blocks (from EUI) projected into the female and male GCA large intestine models.

### Millitome Tissue Registration

Millitomes^27–29^ provide guidance on how to partition an organ, often a complete organ. They are implemented either as a 3-dimensional mold that physically holds an organ for partitioning, or a drawing of an organ that is used as a guide by surgeons when they partition an organ free hand, or both. All tissue blocks cut using millitomes can be automatically registered within HRA 3D reference objects and can be examined in the Exploration User Interface.

To integrate millitome-dissected tissue blocks into the HRA, the digital 3D millitome model must be computationally aligned with the corresponding HRA reference organ. Similar to cross-atlas alignment, AMAP is used for whole organ alignment and tissue block projection. Each millitome can have hundreds of blocks (extraction sites) and all blocks get registered into the HRA and displayed in the EUI, thereby avoiding the time-consuming RUI, and automating and scaling HRA registrations.

Each mesh object is assigned a unique Millitome ID, and blocks without a valid label are removed to eliminate artifacts (see **Methods**). AMAP then generates two key output files: *extraction-sites.jsonld*, encoding the spatial positions and geometries of each tissue block aligned to the reference organ, and *dataset-graph.jsonld*, containing donor and tissue block metadata linked to the extraction sites. These are reviewed and ingested into the HRA Knowledge Graph (HRA-KG), where they become accessible via the Linked Open Data endpoint and the HuBMAP API. To associate experimental datasets, a lookup table (*.CSV file) associates each Millitome ID to a tissue Sample ID, and the RUI Locations Processor produces a final *rui_locations.jsonld* submitted to the HRA registration repository—enabling interactive visualization of mapped blocks and associated data in the EUI.

To exemplify and validate the approach, we worked with four data providers to create 7 millitome models (both sexes and lateralities, where applicable) for five organs (fallopian tube, ovary, uterus, kidney, and pancreas). **Table 1** provides a listing of all millitomes. AMAP-enabled millitome-based registration allowed registration of 727 tissue blocks containing 2,055 datasets from HuBMAP. **Figure 6** shows the millitomes and their transformed models.

**Figure 6.**
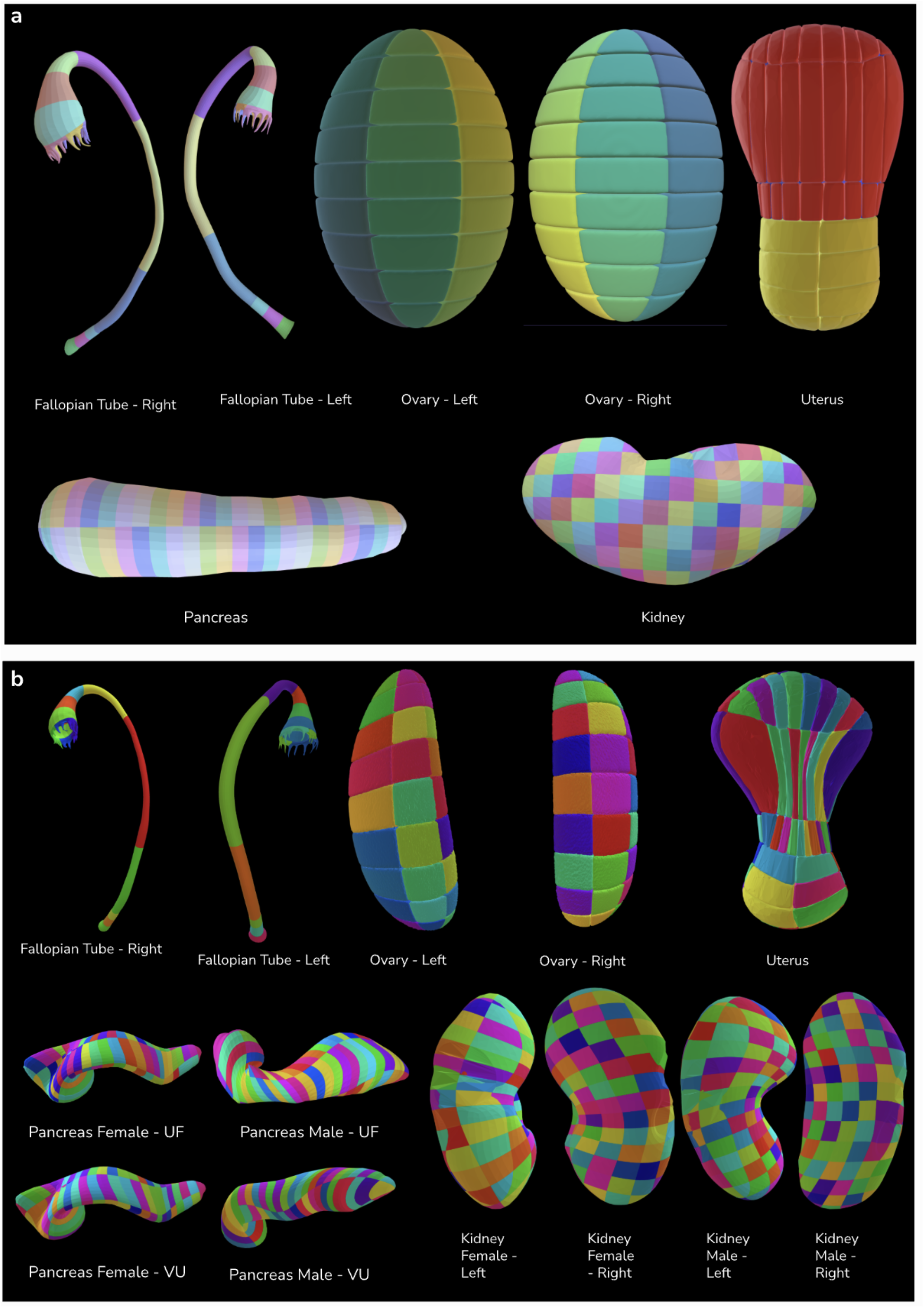
Millitome Tissue Registration. **a.** Shows seven millitome models (virtual 3D GLB models) for fallopian tube (left and right), ovaries (left and right), uterus, pancreas, and kidney. Each model is composed of many individual blocks (extraction sites) such that each block has its own geometry and label (millitome block ID) within the GLB file. The millitome block ID is used in the Lookup Tables for associating extraction sites to datasets. **b.** Shows all millitome models after they are transformed to match the geometries of HRA 3D reference models. The pancreas millitome is a generic pancreas millitome and is mapped to both male and female HRA pancreas models. It is used by different data providers such that the model shape is the same but there are differences in block labeling. Similarly, the kidney millitome model is a generic kidney model and is used to map data to four different HRA kidney models (male/female, left/right). When the millitome models are transformed to match the HRA model geometries, the underlying blocks are also transformed. These transformed blocks are then projected into the HRA EUI.

**Table 1:** A listing of all millitome models used for tissue block registration. Generic kidney model is used to register tissue blocks in male/female kidneys for both lateralities. Generic pancreas model was used to register tissue blocks for both male and female pancreas; both VU and UF use the same pancreas millitome model but with a different number of blocks sectioned.

| Organ | Author | Sex | Laterality | #Blocks |
| --- | --- | --- | --- | --- |
| Fallopian Tube | University of Pennsylvania | Female | Left | 12 |
| Fallopian Tube | University of Pennsylvania | Female | Right | 12 |
| Ovary | University of Pennsylvania | Female | Left | 36 |
| Ovary | University of Pennsylvania | Female | Right | 36 |
| Uterus | University of Pennsylvania | Female | N/A | 132 |
| Kidney | Vanderbilt University | Generic | Generic | 179 |
| Pancreas | University of Florida | Generic | N/A | 121 |
| Pancreas | Vanderbilt University | Generic | N/A | 97 |

## Discussion

The HRA is a multiscale, multimodal, community-driven atlas that integrates diverse assay types across spatial scales—from whole body to single cell—spatially and semantically registering human tissue from 65 organs into a common coordinate framework (CCF) linked to established ontologies. Many other atlasing efforts have focused on different organ systems and different problems of interest, creating their own references and coordinate systems and ingesting data into those atlases. Development of methods that enable better data sharing between different atlas ecosystems^31,32,38^ can help researchers use more comprehensive datasets to solve biomedical problems.

The work presented in this manuscript, makes it possible to interconnect different atlasing systems and lab-specific tissue acquisition systems at the level of spatial and semantic registrations of acquired tissue specimen data. This, in turn, can enable atlases to use their own data and tools in conjunction with those of other atlases. Additionally, it can allow scalable automated registration of future data acquisition into the HRA via use of the millitomes. The work presented here makes it possible to align models from four different atlasing systems as well as 13 millitome models from five different organs.

Several limitations remain: (1) The accuracy of both the whole organ alignment and the block projection relies on visual validation using overlaid point clouds and heatmaps, (2) Since the alignment is computed at the whole organ level, there may be local areas, or anatomical structures, of the models that do not align fully to both the surface and internal geometry, (3) Since the alignment is computed based on mesh surfaces, the internal structures of both the source and target models need to be matched, i.e., if either the source or the target model contain additional structures absent in the other, that particular structure must be removed from the mesh prior to alignment else the source structure may map randomly to any inaccurate target structure to optimize point cloud overlap, (4) The accuracy of alignment is impacted by the difference in point cloud density between the source and target models. Despite these limitations, the AMAP registration pipeline is considered to be “fit for purpose” to exchange data across atlas systems. In fact, tissue registration into reference organs is assumed to have higher error rates given human diversity. For analysis relying on highly accurate registration between two models, a more accurate local registration should be computed, using AMAP registration as an initial starting point. **Figure 7** shows the registration error using Signed Distance Field (see **Methods**) for all models used in this manuscript.

**Figure 7.**
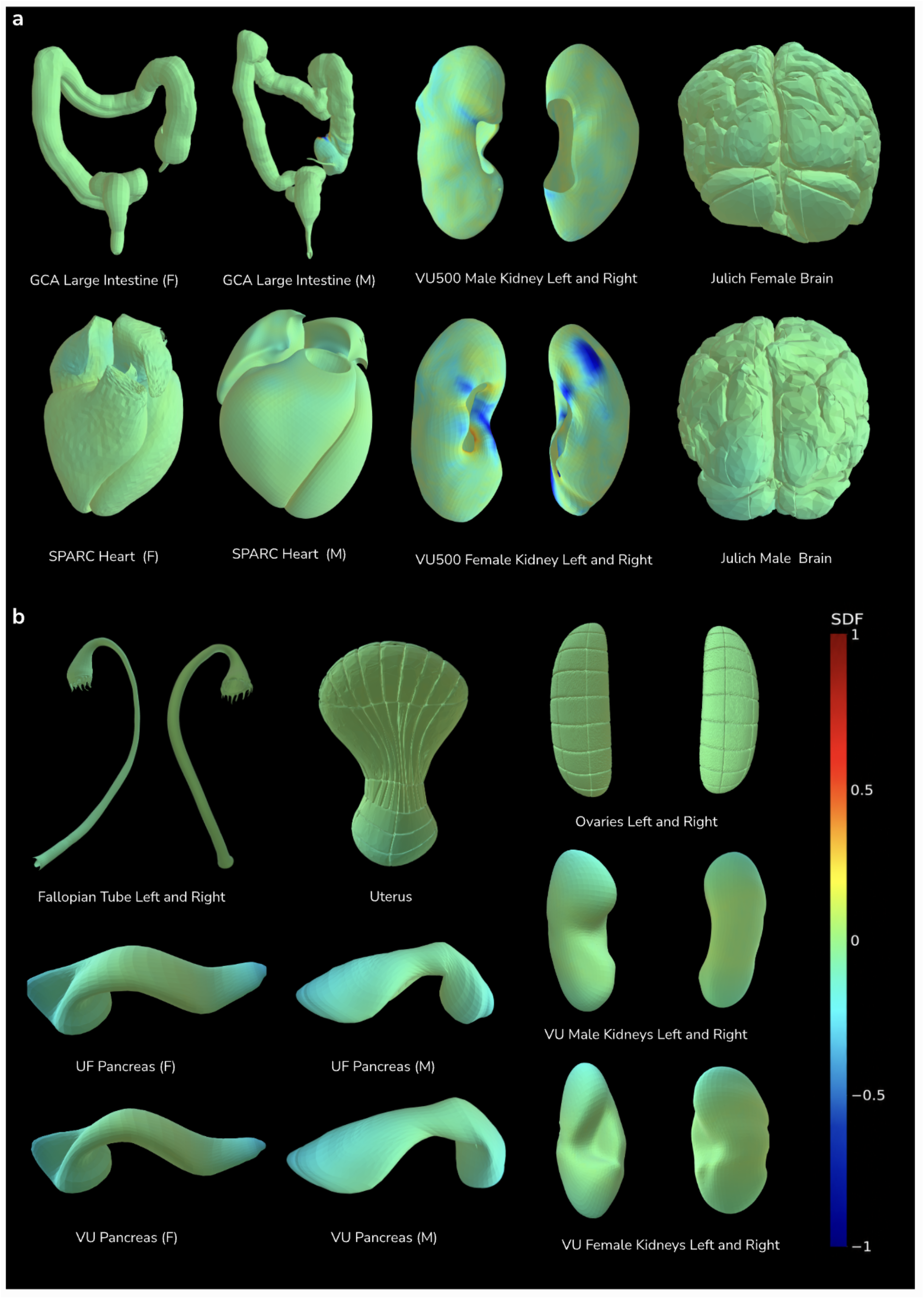
Transformation error visualized as signed distance fields. **a.** non-HRA organs **b.** millitomes. Colors indicate signed surface distance between the transformed source and target anatomy, with green denoting near-zero error, red positive displacement, and blue negative displacement. Signed distances are shown on a normalized spatial scale from -1 to 1.

In the future, we aim to resolve these limitations using approaches such as, but not limited to, registration informed by anatomical structures for better local alignment of features, multi-scale coarse-to-fine registration^39–41^, and hyperparameter grid search. Additionally, we aim to add more atlasing and millitome models to the AMAP library to increase interoperability and reduce data silos across the atlas ecosystems, enabling reproducible, multi-atlas analyses that are not possible within any single reference atlas system.

## Methods

### HRA Reference System

The 3D HRA Reference System (also called common coordinate framework or CCF) is hierarchically nested. **Figure 1** shows the HRA Reference System on the left with the Cartesian coordinate system origin in the back, bottom, left corner. The X-axis of this coordinate system runs from left-to-right, Y-axis as down-to-up and Z-axis as back-to-front as convention. The male and female bodies are placed in this coordinate system, each with their own coordinate system that has the origin in the back, bottom, left corner. Nested organs are placed inside the body. Smaller anatomical structures are placed within organs. Ultimately, cells are placed inside anatomical structures. All 3D objects have an enclosing (8-coordinate) axis-aligned bounding box. The HRA v2.2 supports 71 human organs (including male/female and left/right versions for many organs) and 1,295 spatially modeled 3D anatomical structures, see https://humanatlas.io/3d-reference-library.

### Spatial and semantic registration of tissue specimen data in HRA

The HRA Registration User Interface (RUI) was designed to spatially place (register) tissue blocks inside the 3D male and female HRA bodies. We refer to this as spatial registration (or RUI-registration) in the HRA and it is different from algorithmic registration methods that are used to compute transformations between a source and a target model. The spatial registrations done using the RUI (or using the millitome-AMAP pipeline) can then be explored in the HRA CCF space using the Exploration User Interface (EUI). Tissue specimen data (tissue blocks) spatially registered into a target organ collide with 3D anatomical structures associated with the organ and the volume of block overlap per anatomical structure is computed^11,42^ in the HRA. Computationally, the origin of the organ is first transformed to the origin of the global HRA reference system at (0, 0, 0). A reverse transformation is applied to make the origin of the organ coincide with the origin of the global reference frame (0, 0, 0). This aligns the local coordinate system of the organ to the global coordinate system. Consequently, the registered tissue block coordinates are defined with respect to the global HRA reference system and not with respect to the coordinate system of the target organ.

### Overview of the AMAP pipeline

AMAP first converts the 3D organ meshes staged for alignment to their point-cloud representation with format-specific conversion routines (see **Data**). For surface-only registration, the point cloud is defined by the 3D coordinates of the mesh vertices. For volumetric registration AMAP augments the mesh vertices with interior control points derived from a voxelized visual hull. The pipeline then optimizes alignment between the point-sets using a combination of two types of registration algorithms:

- Rigid registration using Random Sample Consensus (RANSAC)^34^ and Iterative Closest Point (ICP)^35^
- Non-rigid registration using Bayesian Formulation of Coherent Point Drift (BCPD++)^36,37^

For an alignment between two point-clouds (*N* × 3) and (*M* × 3), the registrations are applied sequentially; rigid registration finds the best overlap between the global shapes, which non-rigid registration then fine-tunes for the best local fit. The result is a 4×4 transformation matrix which defines the linear transformation (scale, rotation, and translation) for the best global fit, and a (*N* × 3) non-linear deformation vector field (DVF) which specifies how each of the N points must move for the best local fit.

### Rigid registration using RANSAC and ICP

Typically, the organ point-clouds from different sources will be of different sizes, in arbitrary orientation with each other, and far apart in a shared coordinate space (see **Figure 2**). The distance-minimization or expectation-maximizations (EM) algorithms (such as ICP, BCPD) are sensitive to this initial state of source and target models^43,44^. Thus, a scaling, rotation, and translation to obtain at least a coarse alignment before initiating registration is necessary. A widely used solution is the RANSAC algorithm to provide the initial alignment for ICP registration^43,45^. Subsequently, ICP can provide the initial alignment for the BCPD++ algorithm.

RANSAC is a fast feature-matching algorithm that finds coarse alignment between source and target point clouds. This is followed by a rigid-body transformation using ICP for a tighter overlap by aligning the size, orientation, and centers of two organ point-clouds. Moreover, organs that have laterality, such as the kidneys, benefit from this stage since any necessary flipping transformation (for example, when registering left kidney to a right kidney) can take place here improving the biological accuracy of the registration.

### Non-Rigid registration using BCPD++

The tight overlap between the point-clouds is heavily constrained by linear-only transformation that is based on scaling, rotating, and translation; there still exist all the local variations in the topology that are not accounted for because the shape is still preserved. To obtain higher overlap between source and target models, it is necessary to relax the shape rigidity and allow local deformations such that an improved fit of the source point cloud onto the target point cloud is found. Even so, these deformations cannot be of an arbitrary nature, which will sacrifice biological validity, but instead direct topology to move coherently. A popular registration algorithm that provides such a property is the Coherent Point Drift algorithm^46^. However, for point clouds exceeding 50,000 (common for organ point clouds), the acceleration possibilities are severely limited, along with the control options^37^.

Bayesian Coherent Point Drift (BCPD) defines a prior distribution for motion coherence and estimates the unobserved parameters given the target and source point clouds. Additionally, it exposes a list of interpretable parameters, for example, to control the inertia and length of motion vectors, that can be tuned individually to achieve the correct fit. The algorithm calculates a deformation vector field to move each point on the source point cloud to the target point cloud, constraining the deformations to be locally coherent.

### Projecting tissue blocks after whole organ alignment

Since the set of all transformations that move a point from source to target are known, projecting a tissue block (defined in software by a set of eight bounding box coordinates) from source to target is reduced to applying the transformations in sequential order. Anatomically, however, tissue blocks are volumetric entities. AMAP therefore densely samples each tissue block into a tissue block point cloud and applies the saved transformations (projection). The densely sampled point cloud of the tissue block may contain points not originally present in the registered source point cloud. For points such as these, the non-rigid transformation (the deformation vector field) is undefined; interpolation using a nearest-neighbor scheme determines their projection path. Registering organs volumetrically strengthens interpolation by providing support neighbors to the points in the interior. To recover the block geometry from the projected cloud of the block, AMAP fits an optimal similarity transform between the source and projected point cloud and applies this transform to the source block mesh, eliminating distortions in the mesh geometry from the non-rigid step (see **Figure 7**).

### Quantifying registration error with a signed distance field (SDF)

To quantify geometric disagreement between the transformed (source) and target (original) organ surfaces, we use an SDF-based surface error, where the target organ surface defines the zero-level set and vertices of the transformed source surface are assigned their signed distance to that target surface (see **Figure 7**). SDFs provide a basis for comparing shapes without requiring explicit point-to-point surface correspondences. If object shapes are transformed correctly, the SDFs of their surfaces in the common coordinate system should match^47^.

### Data Formats

3D assets from different sources and providers vary between 3D volumetric image representations (NIfTI) or surface-based 3D anatomical models (GLB, STL). Many popular software libraries exist for image registration of 2D and 3D images that deal with data formats such as NIfTI and DICOM for CT and MRI datasets^48–53^. While such methods can be extended for mesh or point cloud registrations, there are fundamental differences in data representation and handling of deformations and topological changes that make such algorithms unsuitable. There is also a key challenge of converting a 3D object (mesh) to a volumetric (voxelized) image which is a challenging and ambiguous transformation task. Surface (mesh or point cloud) registration algorithms^35,54,55^ are designed to align and deform object meshes (composed of vertices and faces) to establish correspondences between different shapes or anatomical structures.

All HRA 3D reference objects are provided as GLB (*.glb) files. Data for non-HRA models and millitomes come in a variety of formats (VTK, STL, FBX, GLB) and were converted as follows:

### Conversion to GLB

All non-GLB models are converted into a triangle mesh structure, i.e., the representation of a 3D object as approximated by triangles to build and represent the surface of that object. The resulting file has at least two types of information, (a) the vertices of a triangle or a face, (b) how the triangles or faces are connected to each other, and (optionally) (c) block labels for each vertex in case of conversion from a millitome model (to distinguish individual millitome blocks apart from the others). These converted objects are exported as GLB files and are made openly available (see **Data Availability**). Trimesh^56^ (https://trimesh.org), a Python library, is used to do the conversions.

### Conversion to point clouds

Next, the GLB files are converted into point clouds using the vertices from the mesh. Note that this approach makes the point cloud representation reversible, such that the original organ object mesh can be uniquely obtained from the point cloud (after running it through the registration engine) by leveraging the face connectivity information in the original mesh structure.

### Final Projections

The final transformation matrices are saved as a serialized Python object (*.pickle.gz file).

### Data

#### HRA 3D Reference Models

The HRA 3D reference object library provides anatomically correct open-source 3D reference organs used for tissue specimen data registration (RUI) and exploration (EUI). This study focuses on a subset of available 3D reference objects in the HRA based on the different use cases elucidated in the **Results** section. All versions of all HRA 3D reference objects can be accessed at https://humanatlas.io/3d-reference-library.

#### Non-HRA Atlas Models

Other atlasing efforts are detailed below.

#### Gut Cell Atlas Large Intestine

The Gut Cell Atlas^23^ reference system abstracts the gut as a 1D centerline from the gastroduodenal junction to the anus, with location encoded as endoscope-measured distance to standardized anatomical landmarks. It has been extended to 2D and 3D using anatomograms and anonymized CT scans, with off-centerline locations resolved by projection to the nearest midline point. In prior work^23^, the GCA team created 3D versions of their 1D models and also created versions transformed and aligned with the HRA large intestine model. We use this specific 3D model in this work.

#### 3D SPARC Heart

The SPARC Program^25^ developed an annotated generic human heart scaffold, derived from average organ dimensions in the literature, as a registration target for experimental datasets on the SPARC Portal, primarily to support neuromodulation device research.

#### 3D Julich Brain Atlas

The Julich Brain Atlas^22^ is a continuously expanding probabilistic 3D brain atlas integrating cytoarchitectonic maps, connectivity data, neurotransmitter receptor profiles, and functional data in a common reference space. It is openly accessible and designed for interoperability with other brain parcellations and databases. We use the official siibra-python^32^ API (https://siibra-python.readthedocs.io/en/latest) to access the 3D brain model and use the model created in the *MNI_COLIN_27* template space.

#### VU500 consensus kidney atlas

The VU500 Consensus Kidney^26^ is constructed from multi-contrast CT scans of 500 healthy subjects and captures kidney morphometry across non-contrast and four contrast-enhanced phases using a two-stage hierarchical registration pipeline combining metric-based and deep learning methods.

### Millitomes

For AMAP-compatibility, each millitome requires a virtual model (3D mesh) where each block is a separately defined geometry in the 3D mesh. These are generally saved as GLB files.

We worked with four data providers to create 7 different millitome models for five organs (see **Table 1** and **Figure 6**). The block specifications for each model (shape, number, size) were defined in collaboration with the data providers according to their needs and sectioning protocols. While some models use the HRA reference models as the base for millitome (fallopian tube, kidney), the others are based on their own versions of 3D models and hence differ in shape from their HRA counterparts (uterus, ovaries, pancreas). Each millitome model has corresponding *Lookup Tables* (CSV file; see **Figure 1 bottom panel**) that maps grid positions to Millitome IDs and includes metadata such as organ type, dimensions, and block size. This table is used to associate sample and dataset metadata to individual extraction sites in the HRA EUI.

Prior Standard Operating Procedures discuss standards and processes for creating and using millitomes^27–29^ (see also, https://github.com/hubmapconsortium/hra-registrations/blob/main/hubmap-fallopian_tube_millitome-fisher-2024/resources/HuBMAP%20Labels%204-3-24.pdf). All final millitomes are published on GitHub (see **Data and Code Availability**) as well as released as part of the HRA Knowledge Graph^57^ with individual DOIs (https://apps.humanatlas.io/kg-explorer/?do=millitome).

Since the models can be based on varied sources, the organ meshes may contain both external and internal anatomical structures. To prepare the models for use in mold construction or AMAP alignment, non-essential internal geometries (e.g., vasculature, inner parenchyma) must be removed so that the source and target models have similar geometries.

This cleaning is performed in 3D content creation software such as Blender. The steps include:

1. **Import Organ Mesh:** The 3D reference organ model is imported into a 3D modeling environment such as Blender or Cinema 4D.
2. **Isolate Outer Shell:** Using the scene hierarchy and visual inspection, the outermost mesh capturing surface boundary of the organ is identified. This is the only structure required for millitome mold construction, as internal components are not needed for defining the cutting geometry.
3. **Remove Internal Structures and Artifacts:** All internal anatomical meshes, nested elements, and non-essential geometries are deleted. Any stray polygons or disconnected fragments are also removed to ensure the resulting mesh is a single, watertight (manifold) shell suitable for Boolean operations.
4. **Center the Mesh Origin:** The mesh origin (pivot point) is relocated to the geometric center, ensuring that the organ is positioned correctly relative to the base of the millitome mold.
5. **Align Mesh to Standard Coordinate Frame:** The cleaned mesh is rotated such that it lies flat along the XY plane, with its longitudinal axis aligned to the X-axis. This standardized orientation ensures consistency across organ models and facilitates uniform cutting grid placement during millitome generation.

In summary, the 3D printed millitome mold is used to dissect fresh organs; tissue blocks are tracked using Millitome IDs and assigned Sample IDs during collection and shared via *CSV Lookup Tables*. This tissue specimen data is submitted into HRA via the virtual 3D millitome model and the AMAP pipeline to support downstream HRA integration and EUI visualization.

## Data Availability

All data (non-HRA atlas models, millitomes) used in the work, in addition to computed transformation matrices, and transformed models are made available via GitHub (https://github.com/cns-iu/hra-amap; see */input-data*, */output-data*, */raw-data* directories).

## Code Availability

All code is made publicly available via GitHub at https://github.com/cns-iu/hra-amap.

## Author Contributions

**YJ** co-led the paper writing and study design, guided AMAP and millitome design decisions; **BD** designed and implemented the core AMAP pipeline, contributed fixes and improvements to the AMAP codebase and aided production, contributed to paper writing; **DQ** worked on millitome registrations and metadata acquisition, paper writing, and study design; **AB** contributed to the AMAP codebase, refactored it for production use, and processed the millitome and non-HRA models for ingestion into HRA-KG; **PK** contributed to the 3D millitome modeling; **AP** contributed to the 3D millitome modeling; **KO** contributed to the 3D millitome modeling; **SA** worked on millitome registrations and metadata acquisition; **BH** led the technical development of the AMAP codebase, implemented HRA-KG, HRA-API, and EUI support for millitomes, and worked with millitome authors to validate results; **SF** worked on millitome generation, registrations and metadata acquisition, and paper revisions; **KB** leads the Human Reference Atlas, conceptualized the work and contributed to writing and revising of manuscript.

## Acknowledgements

We thank all the data providers who collaborated with us to create and use the millitome models: Vanderbilt University (Jeff Spraggins, Jamie Allen, Diane Saunders), University of Florida (Martha Campbell-Thompson).

This work was funded by the NIH Common Fund through the Office of Strategic Coordination/Office of the NIH Director under awards OT2OD033756, OT2OD026671, OT2OD030545, and R03OD039970, by the Cellular Senescence Network (SenNet) Consortium through the Consortium Organization and Data Coordinating Center (CODCC) under award number U24CA268108 and via award U54AG076043, and by the NIDDK under awards U24DK135157 and U01DK133090. The funders had no role in study design, data collection and analysis, decision to publish, or preparation of the manuscript. The content is solely the responsibility of the authors and does not necessarily represent the official views of the National Institutes of Health.

